## Supplementary Materials for "Imaging fascicular organization of peripheral nerves with fast neural Electrical Impedance Tomography (EIT)"

#### Neural tracing and effects on evoked compound action potentials in sciatic nerve

In several preliminary experiments, we attempted neural tracing in rat sciatic nerve using three types of conventional tracers – FluoroGold, DiI, fluorescent dextran conjugates, as well as five different adeno-associated viral vectors. The viral vectors driving the expression of fluorescent reporter gene under control of neural-specific promoter were injected intramuscularly into gastrocnemius medial and lateral muscles, or into tibialis anterior muscle of 8-week old rats: 25 µL suspension containing $5\text{-}10\times10^{11}$ vg was delivered across three sites per muscle using a 30G needle at 5 µL/min as described in[1]. Viruses used in our study: AAV2-hSyn-eGFP (UNC Vector Core), AAV5-hSyn-GFP (UNC Vector Core), AAV5-hSyn-eYFP (UNC Vector Core), AAV9-hSyn-eGFP (Penn Vector Core), and AAV2-CamKII-tdTomato (custom made). However, 5-6 weeks after injection, no axonal labelling at the level of sciatic nerve was observed. The plausible explanations are that the AAVs are generally not efficient when used for retrograde tracing, and a relatively low titre of the viruses used.

The fluorescent tracers FluoroGold (FG) and DiI were injected intraneurally into tibial or peroneal branches of the sciatic nerve using glass micropipette attached to micromanipulator at 0.5-1 µL/min (2 µl of 4% FG in saline or 3 µl of 5% DiI in 10% DMSO). 24-48h post-injection, the axonal labelling at the level of common sciatic nerve was achieved. However, both tracers caused significant impairment to motor function. We, therefore, opted for non-toxic dextran-based neural tracers – fluorescent dextran conjugates Alexa Fluor 488 and Alexa Fluor 555 dextran 10 kDa (ThermoFisher). The speed of tracer diffusion within the nerve was evaluated to allow for optimal staining of the fascicles at the level of the common sciatic nerve at the time of the EIT experiment and subsequent histological and microCT analysis. The different time points tested were as follows: 6h, 12h, 24h, 48h, 72h and 120h. An incubation time of 48h was chosen as optimal – this period allows sufficient diffusion of the tracer to the level of common sciatic nerve when injected into both peroneal and tibial fascicles. Shorter periods (6-24h) resulted in diffusion up to the branching region of the nerve only (~0.5-1 cm distally from the area where the EIT cuff electrode is placed during the experiment), while longer period (72-120h) resulted in insufficient staining of the fascicles at the level of the EIT cuff electrode placement due to diffusion of the tracer to the more proximal region of the nerve. The diffusion speed was ~2 cm/day and is in accordance with previously published studies[2]. Resulting fluorescent images after 48h incubation time were of high contrast and specificity, with clearly distinguishable boundaries of the fascicles (peroneal and tibial) at the level of EIT cuff placement (Supp. Fig. 1a).

It is known that fluorescent neural tracers can affect the nerve electrophysiology[3]. Additionally, the procedure of tracer application can be traumatic due to local volume/pressure perturbations during tracer injection. Indeed, we observed that FG and DiI cause significant impairment of the nerve electrophysiology, at least in a time frame of our experiments (1-5d post-injection). In contrast, fluorescent conjugates of dextran caused no abnormalities of motor function in any of the animals included in the study. In one animal, tracer was injected into the peroneal branch of the left sciatic nerve (to check the effect of the tracer on evoked CAP magnitude), while a control nerve received injection of the same volume of saline. After 48h, the effects of the tracer and saline injections were evaluated using stimulation of the peroneal branch in the left (injected with tracer) and right leg (control) (Supp. Fig. 1b). The magnitude of the recorded CAPs (400-600 µV, Supp. Fig. 1b) was not significantly affected by tracer injection and was within the range observed in EIT experiments without neural tracers[4].

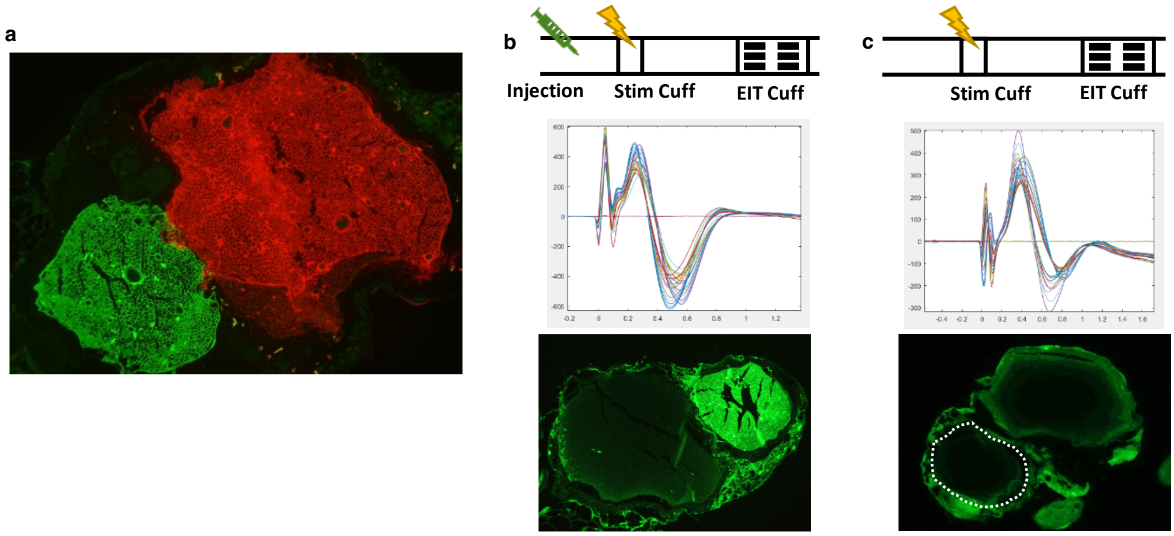


**Supplementary Fig 1. Neural tracing and its effect on CAP**. **a**, Representative example of labelling of the tibial (red) and peroneal (green) fascicles with fluorescent dextran conjugates at the level of EIT cuff placement. **a**, Effects of neural tracer injection on magnitude of evoked CAPs in the peroneal branch (left) as compared to CAPs in a control nerve (right, the white dashed line demarcates peroneal branch without tracer).

### Cross-validation of techniques and CoM localization errors

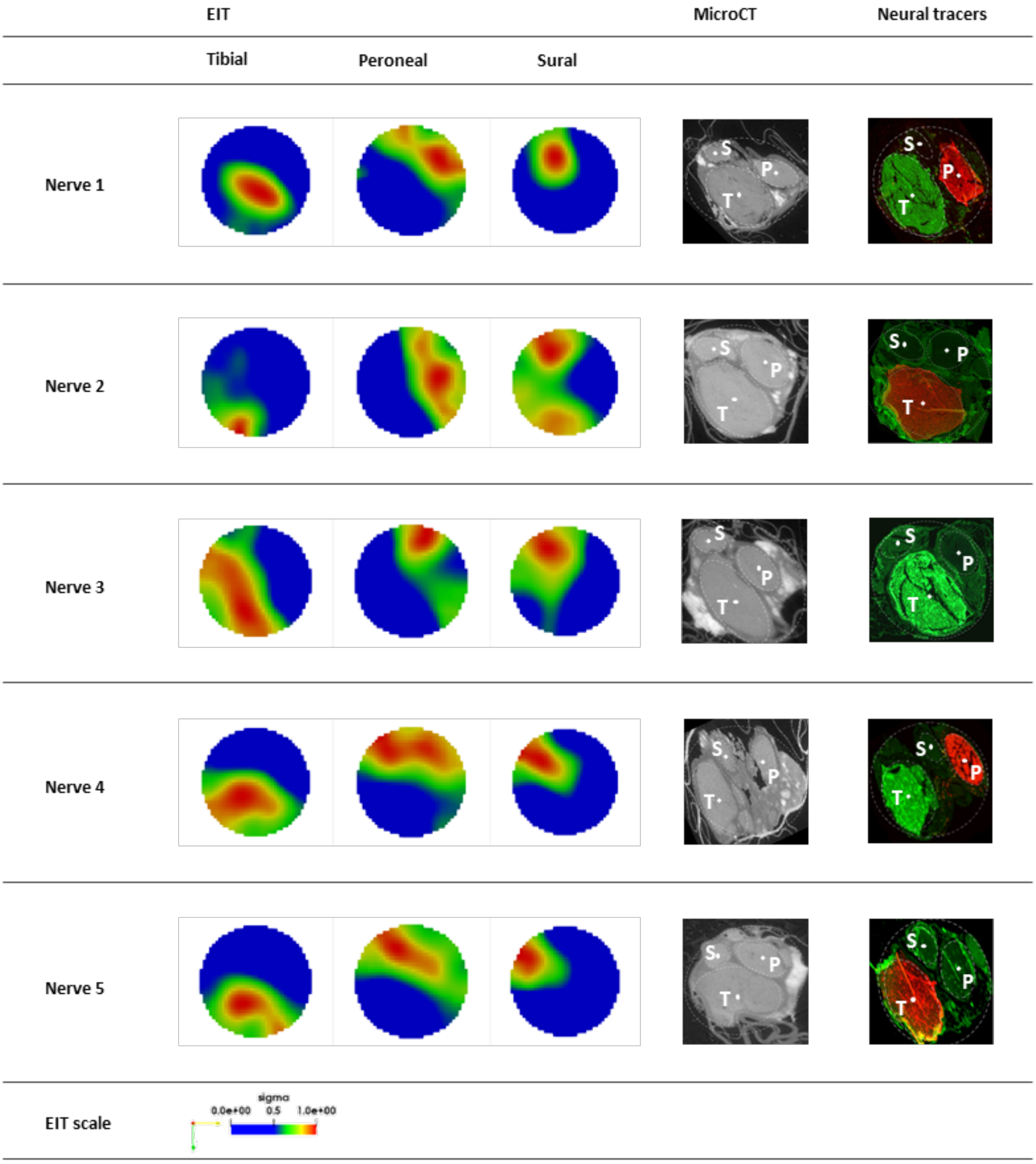


**Supplementary Fig 2. Co-registered images of all five nerves for EIT, microCT and neural tracers**. *EIT:* the individual images of evoked fascicular activity reconstructed from EIT recordings for the tibial, peroneal and sural fascicles, respectively. The range of values for every image was normalized between 0 and 1 and the rainbow color scale is showing the top 50% of color for each image i.e. full width at half maximum (FWHM) intensity scaling. *MicroCT:* representative XY plane slice of the microCT scan of the nerve in the area corresponding to EIT recordings with fascicles clearly visible and snippets of the silk thread marking the EIT cuff opening area. *Neural tracers:* The fluorescent image of the cross-section of the common sciatic nerve at the area corresponding to EIT recordings with tibial and peroneal fascicle labelled with either fluorescent Dextran-Alexa 488 (green) or Dextran-Alexa 555 (red). The margins of the sural fascicle which did not receive tracer injection are demarcated with a dashed line. Each microCT and neural tracer image has undergone rigid deformation and rotation for purposes of cross-validation of CoM of fascicles between three techniques. All images were rotated so that the electrode cuff opening was at 12 o'clock. The external boundaries of the nerve were fitted to a circular profile after rigid deformation. Each fascicle was fitted into an ellipsoid profile (dashed white line) with a CoM (white dot in the middle) which was further used as a ground truth coordinate in the cross-validation study.

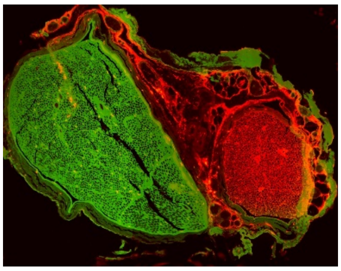

**Supplementary Fig 3. Fluorescent tracers for Nerve 3**. In Supplementary Fig. 2, the neural tracer image for Nerve 3 did not contain evidence of Dextran-Alexa 555 (red) in the peroneal fascicle at the position in the common sciatic nerve corresponding to EIT and where all three fascicles were visible. However, another cross-section image of the nerve, shown here, where only the tibial and peroneal fascicles were visible, shows the presence of both neural tracers. This, along with the corresponding microCT image, allowed for the inference of the position of the peroneal fascicle in the neural tracer image shown in Supplementary Fig. 2.

**Supplementary Table 1. Comparison of three techniques for their ability to discriminate radial (R) and angular (Theta) position of individual fascicles (tibial, peroneal and sural) within the common sciatic nerve (15 fascicles in N=5 nerves)**

| Fascicle | EIT | | MicroCT | | Neural tracers | |
|---|---|---|---|---|---|---|
| | R [µm] | ϑ[°] | R [µm] | ϑ [°] | R [µm] | ϑ [°] |
| Tibial (T) | 272±57 | -124±27 | 255±79 | -119±14 | 270±55 | -126±24 |
| Sural (S) | 287±86 | 145±24 | 501±76 | 131±4 | 482±71 | 112±15 |
| Peroneal (P) | 275±38 | 49±22 | 345±73 | 39±13 | 407±72 | 35±17 |

**Supplementary Table 2. CoM localization errors between EIT, microCT and neural tracer histology**
(Errors are quantified as normalized distance vector)

| Experiment | Fascicle | Error EIT vs µCT, µm | Error EIT vs histology, µm | Error µCT vs histology, µm |
|---|---|---|---|---|
| Nerve 1 | T | 61 | 220 | 166 |
| | S | 241 | 188 | 251 |
| | P | 268 | 221 | 103 |
| Nerve 2 | T | 182 | 250 | 72 |
| | S | 446 | 444 | 95 |
| | P | 124 | 271 | 152 |
| Nerve 3 | T | 181 | 197 | 35 |
| | S | 337 | 332 | 64 |
| | P | 50 | 170 | 201 |
| Nerve 4 | T | 213 | 162 | 77 |
| | S | 94 | 277 | 237 |
| | P | 134 | 339 | 205 |
| Nerve 5 | T | 122 | 208 | 157 |
| | S | 95 | 282 | 244 |
| | P | 146 | 204 | 127 |
| | | **178±108.1** | **251±76.2** | **146±70.9** |
